# Analysis of cell size homeostasis at the single-cell and population level

**DOI:** 10.1101/338632

**Authors:** Philipp Thomas

**Affiliations:** Department of Mathematics, Imperial College London, London SW7 2AZ, UK

**Author notes:** Electronic mail.

## Abstract

Growth pervades all areas of life from single cells to cell populations to tissues. However, cell size often fluctuates significantly from cell to cell and from generation to generation. Here we present a unified framework to predict the statistics of cell size variations within a lineage tree of a proliferating population. We analytically characterise (i) the distributions of cell size snapshots, (ii) the distribution within a population tree, and (iii) the distribution of lineages across the tree. Surprisingly, these size distributions differ significantly from observing single cells in isolation. In populations, cells seemingly grow to different sizes, typically exhibit less cell-to-cell variability and often display qualitatively different sensitivities to cell cycle noise and division errors. We demonstrate the key findings using recent single-cell data and elaborate on the implications for the ability of cells to maintain a narrow size distribution and the emergence of different power laws in these distributions.

## I. INTRODUCTION

Cells decide when to divide based on their size. A key question is therefore why cells grow to a certain size, how they maintain these sizes within a narrow distribution and what are the dominant sources of size variations. Recent experiments investigate this issue by taking snapshots of populations^1^, trapping single cells^2^ or recording movies of growing populations^3?^. Currently, we lack the tools with which to compare these different data.

It is known that timing divisions independently of cell size, a so-called timer control, does not lead to a stable size distribution when cells grow exponentially^4,5^. Since exponential growth is ubiquitously observed in microbes^2,3,6-8^, cells must control their division sizes to achieve homeostasis^9?,10^. A possible strategy for such a control, called a sizer, is to set a stochastic threshold. Many microbes, however, rather grow by a constant size from birth to division, called the adder control^3,11-13^. Other mixed strategies may be described by sizer-or timer-like controls^8,13,14^.

Identifying which strategy cells employ is essential to understand how cells compensate for errors in size control and cell division. Early theory focussed on the distribution of cell size snapshots across a growing population, mainly due to the experimental limitations of tracking many cells over time^15-18^. These initial studies formulated sizer models based on the population balance equation, an integro-differential equation that is notoriously difficult to solve in practice^19-21^. With the advent of single-cell traps such as the mother machine^2^, the theoretical focus moved towards describing single cells over many divisions^11,22-24^. This path is more amenable to analysis because it considers only a single individual described by a discrete-time stochastic process or stochastic map^25^. It does not, however, apply to time-lapse movies monitoring cell growth in populations. A quantitative investigation of this issue is essential for our understanding of cellular physiology.

Here, we develop a comprehensive quantitative framework to predict single-cell and population statistics from models of cell size control. We focus on a stochastic branching process that produces a lineage tree (Fig. 1a). We analyse (i) the size distributions of single dividing cells followed over time, (ii) the distribution in lineages across a population tree, (iii) the distribution across all cells in the tree, and (iv) the distributions of snapshots. Although these measures stem from the same lineage tree and the same underlying stochastic process, we find that they can give quantitatively and qualitatively different results when size varies from cell to cell. Our findings highlight the significance of population dynamics for the analysis of cell size control and size homeostasis.

**FIG. 1.**
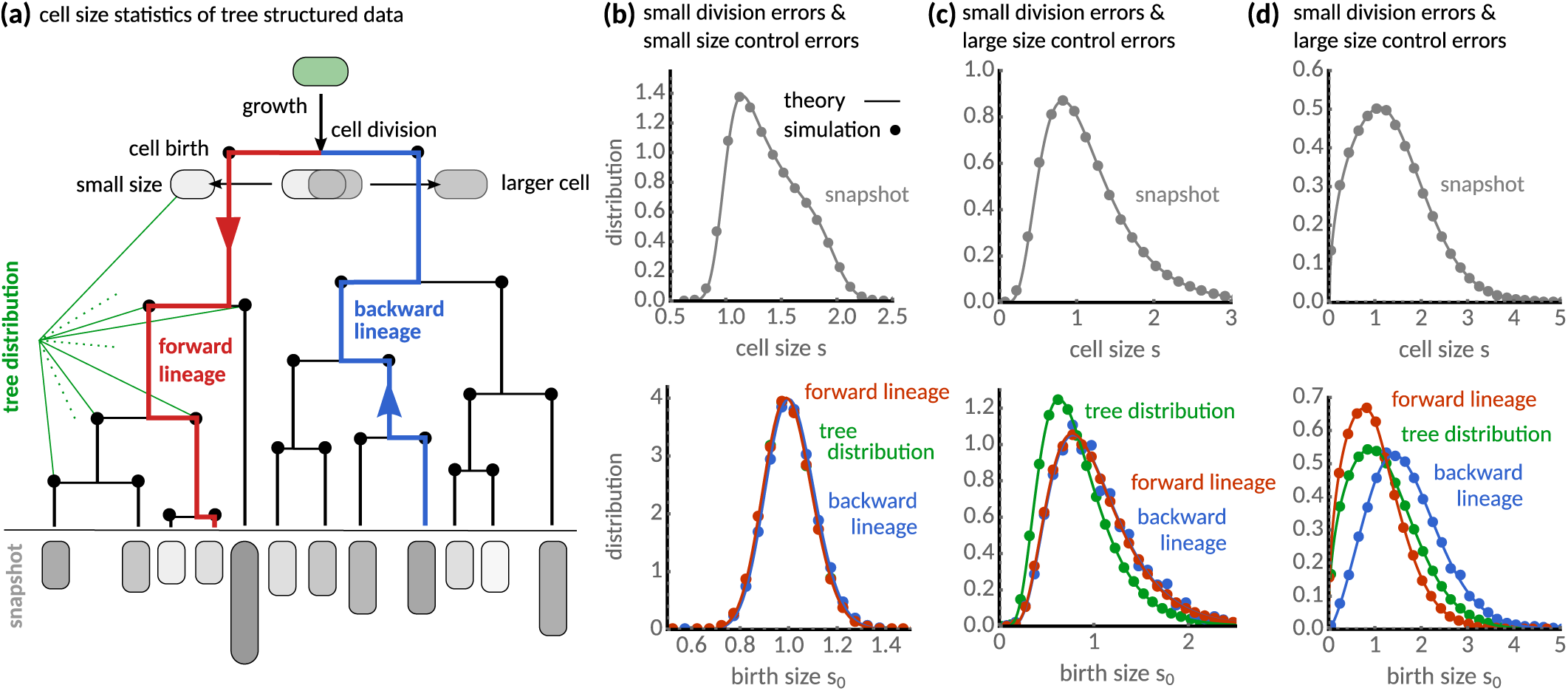
Characterisation of cell size statistics from tree-structured data. **(a)** Lineage tree resulting from cell growth and division. The snapshot distribution characterises the final state of the tree (grey). The distribution of birth sizes resulting from all cell divisions in the tree (black nodes) is the tree distribution. Forward lineages (red lineage) traverse the tree from root to leaf and follow randomly each daughter cell after division. Backward lineages start a randomly chosen cell in the final population and hence traverse the tree from leaf to root. **(b)** The snapshot distribution (grey) for the case of small division errors shows a maximum for newborn cells and decreases progressively towards larger cell sizes. Forward (red), backward lineage (blue) and tree distributions (green) of birth size at birth are qualitatively similar. **(c)** For large fluctuations in cell size controls (CV_*η*_ = 0.75) the snapshot distribution broadens significantly (grey). Distributions of forward and backward lineages are almost indistinguishable, while the tree distribution (green) shows that cells are typically smaller compared to the lineage statistics. **(d)** For large division errors (CV_*p*_ = 0.5), forward, backward lineage and tree distributions appear all significantly different showing cells in forward lineages are typically smaller than in backward and tree distribution. The analytical results (solid lines) are verified using stochastic simulations in all cases (dots). Theory and simulations use adder rule (*a* = 1), gamma-distributed cell cycle noise with *E*[*η*] = 1 and CV_*η*_ =0.1, and division errors following a symmetric beta distribution with CV_*p*_ = 0.07 unless stated otherwise.

## II. ESTIMATION OF CELL SIZE DISTRIBUTIONS FROM TREE-STRUCTURED DATA

Estimation of cell size from snapshots samples all cells at a given instance in time. Time-lapse microscopy, however, offers various statistical analyses based on the underlying tree-structure of the data. A tree corresponds to an experiment starting from a single cell that resolves all ancestral relationships between cells in the population (see Fig. 1a). A lineage is a subset of the tree representing a line of descent between the ancestral cell (root of the tree) and a cell of the final population (leave).

To estimate the tree and lineage statistics, we use the lineage counting method proposed by^26^. We denote by 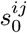 the size measurements at cell birth of the *i^th^* of *D_j_* cells in the *j^th^* of *N_L_* lineages of *N_T_* trees and by *D_j_* the number of cell divisions in the *j^th^* lineage. There are several ways in which these data can be pooled to represent either tree or lineage statistics. The corresponding distributions and their moments therefore generally represent weighted averages of the form

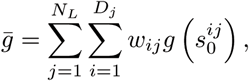

where *w*_*ij*_ are the weights and the function *g* denotes the desired summary statistic. Choosing, for instance, *g* to be the Dirac delta function (or an appropriate binning function) gives a weighted cell size distribution. Similarly, choosing *g*(*x*) = *x*^*n*^ we estimate its *n*^*th*^ moment. In this article, we consider the distributions that correspond to the following weights:

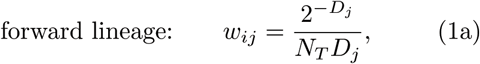

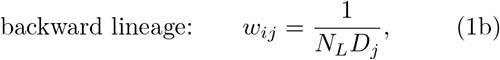

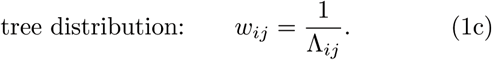

The *forward lineage* is a random path in a tree traversed from root to leaves in which each daughter cell is chosen with equal probability. The weights (1a) follow from the fact that the probability of choosing such a lineage from *N_T_* binary trees is 2^−*D*_*j*_^ / *N_T_* and there are *D_j_* measurements in the *j^th^* lineage (see^8,26^ for details). A similar lineage would be observed in the mother machine^2^ assuming mother and daughter cells are statistically indistinguishable.

The *backward lineage* gives equal weight to each lineage and thus characterises a typical lineage in a tree. The weights (1b) follows because there are *D*_*j*_ measurements in the *j*^*th*^ lineage and *N*_*L*_ lineages. Such a lineage can be thought of choosing a random cell of the final population and traversing the tree from leave to root as described in ^26-28^.

Finally, the *tree distribution* gives equal weight to each birth event in the tree. Consequently, the factor Λ_*ij*_ denotes the number of occurrences of the values 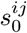 in the *N*_*L*_ lineages. Note that in practice this measure is computed easily by averaging all birth sizes in the trees. In the following, we develop an analytical framework to obtain these distributions from models of cell size control.

## III. SIZE DISTRIBUTION OF A SINGLE CELL OVER TIME

As explained in the previous section, the forward lineage samples either daughter cell with equal probability. We can model such paths as stochastic maps with respect to the number of cell divisions *D*^25^. Denoting the birth size 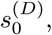 division size 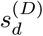 and inherited size fraction *p*^(*D*)^, we have

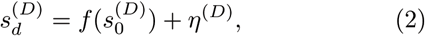

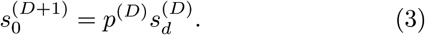

Here the deterministic function *ƒ* determines how division size depends on birth size and the stochastic part *η*^(*D*)^ is independent of birth size. An equivalent formulation is the one in terms of probabilities. In the limit of long times, the birth size distribution 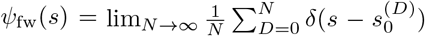 is stationary and satisfies the integral equation

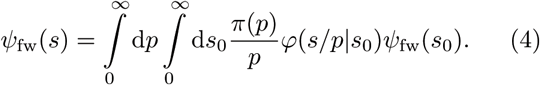

The distribution *φ*(*s|s*_0_) = *φ*_*η*_(*s*_*d*_ − *ƒ*(*s*_0_)) can be written in terms of the distribution of the birth-size independent part of division size *η* and *π*(*p*) is the distribution of inherited size fractions *p*. Because we follow each daughter cell with equal probability, the latter can be written in terms of the marginal distributions of the two daughter cells

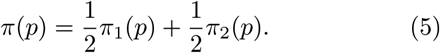

Because the mother’s size is split between the daughters, we have *π*_1_(*p*) = *π*_2_(1 − *p*) which implies that *E*[*p*] = 1/2 and also models the situations of asymmetric cell divisions. Hence on average a daughter cell inherits precisely half of the mother’s size, as expected.

## IV. SIZE DISTRIBUTIONS IN GROWING CELL POPULATIONS

While the theory of a single dividing cell is well established, individual-based frameworks of growing cell populations are less commonly considered but date back to^15^. We go on to develop this theory by deriving the governing equations for the population statistics. Assuming that cell division occurs at a rate *γ*(*τ, s*) that depends on the present cell age *τ* and size *s*, the rate of change in the number of cells with that age and size *n*(*τ, s*) is

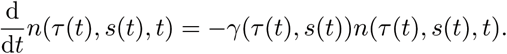

We further assume that cells grow exponentially with growth rate *α*

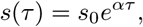

as has been reported for many microbes such as *E. coli*^2^, *C. crescentus*^3^, *B. subtilis*^6^, *M. smegmatis*^8^ and *S. cerevisiae*^7^. Using this dependence we can expand the total derivative to obtain

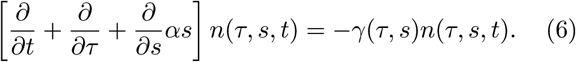

To account for cell births the cell density must obey the boundary condition

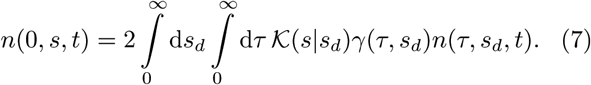

The condition implements the fact that the number of newborn cells in the population must equal twice the number of dividing cells. The division kernel 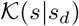 denotes the transition probability for a cell of size *s*_*d*_ to inherit size *s* after cell division. Most conveniently, we can write it in terms of the inherited size fraction via

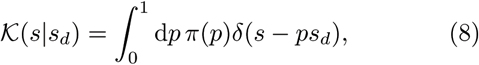

where *π*(*p*) is defined as in Eq. (5) and also takes into account a possible asymmetry between the daughter cells. Similar formulations have been used to characterise distributions of size-structured populations^19,21,29^. In the following, we derive a strategy of how to obtain the various population statistics analytically. We will do this iteratively by writing the snapshot distribution in terms of the tree distribution, and writing the tree distribution in terms of the backward lineage distribution that can be efficiently solved.

### A. Snapshot distribution

In balanced growth conditions the total number of cells increases exponentially, a condition which dramatically simplifies the analysis. In this limit, the cell density follows

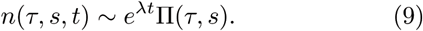

The distribution ∏(*τ, s*) is the probability density of observing a cell with age *τ* and size *s* in a snapshot, which is independent of time. It can further be shown that the asymptotic population growth rate *λ* equals the rate of cell-growth *α*^28^. Using this ansatz in Eq. (6) and employing the method of characteristics, we find that the snapshot density satisfies

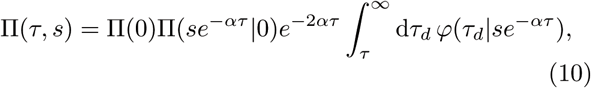

where ∏(*s*_0_|0) is the distribution of birth sizes and

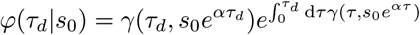

is the distribution of division times for a cell of given birth size. Integrating Eq. (6) and using the boundary condition (7) shows that ∏(0) = 2*α*.

To make the dependence on birth size explicit, we change variables from *τ* to *s*_0_. The result is the snapshot distribution of cell size and birth size, which reads

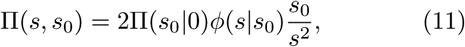

where 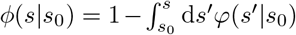 is the probability that a cell born with size *s*_0_ has not divided before reaching size *s*. The snapshot distribution (11) depends explicitly on the distribution of birth sizes, which we characterise in the following. In the absence of cell division errors and cell cycle noise, the birth size *s*_0_ is deterministic and the snapshot distribution reduces to ∏(*s*) = 2*s*_0_/*s*^2^ for *s*_0_ ≤ *s* ≤ 2*s*_0_ as shown by^15^.

### B. Tree distribution

In balanced growth conditions, the fraction of cells with a certain size is constant. Hence, the birth size distribution in a snapshot is also the distribution of birth sizes in the population tree

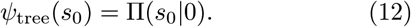

To characterise this measure, we substitute Eqs. (9) and (10) into the boundary condition (7) to find

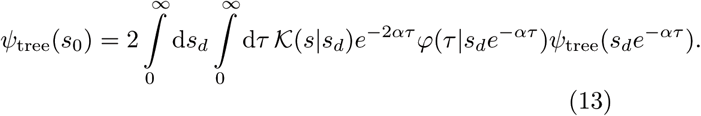

Changing the integration from *τ* to *s*_0_ = *e*^−*α τ*^*s*_*d*_ and using 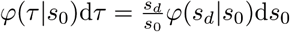, we finally arrive at

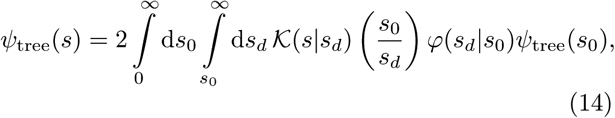

which characterises the tree distribution. The distribution *φ*(*s*_*d*_|*s*_0_) is the distribution of division sizes that describes the cell size control as in Eq. (4).

### C. Backward lineage distribution

The equation for the tree distribution has no intuitive stochastic process interpretation. However, we notice that substituting

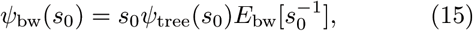

where 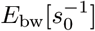 is a normalising factor, transforms Eq. (14) into

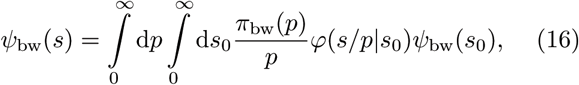

with

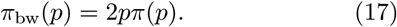

This equation provides such a simple stochastic interpretation because it is of the same form as the forward lineage equation (4), except that *π*_bw_ replaces *π*. In fact, it describes the distribution of birth sizes in a backward lineage. Interestingly, the inherited size fraction in backward lineages *π*_*bw*_ is skewed towards cells inheriting larger size fractions. It thus appears as if cells are dividing asymmetrically in backward lineages. The impact of this phenomenon on the cell size distributions is explored in the following.

### D. Comparison of cell size distributions

A phenomenological linear relation between birth and division size has been observed in experiments^8,13,25^

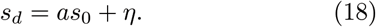

To incorporate these dependencies into our stochastic framework, we set *φ*(*s*_*d*_|*s*_0_) = *φ*_*η*_(*s*_*d*_ − *as*_0_). The parameter *a* denotes different models of cell size control and *φ*_*η*_ the size control distribution. For *a* = 0 division size is independent of birth size, called the sizer mechanism. For *a* = 1 the size added from birth to division varies independently of birth size, an adder mechanism that is commonly observed in bacteria^3,11?^. For *a* = 2 division size is directly proportional to birth size, which resembles cell-size control based on division timing. Intermediate values would behave either sizer-(0 < *a* < 1) or timer-like (1 < *a* < 2).

Eq. (16) could in principle be solved by sampling the stochastic maps given in Sec. III. More effective, however, is to discretise it and solve the resulting set of linear equations numerically. This procedure can in principle be used for arbitrary distributions of division errors and size control. In Fig. 1b,c and d we show the result of this procedure for beta-distributed division errors and gamma-distributed cell size control. The snapshot distribution is obtained using the numerical result for *ψ*_bw_ together with Eq. (16) in (11).

When errors in cell division and size control are small (Fig. 1b), the snapshot distribution displays a maximum at newborn sizes and a long decaying shoulder towards the division size consistent with the fact that in populations the number of newborn cells must equal twice the number of dividing cells. The numerical results for the backward distribution *ψ*_bw_ shown in Fig. 1b are qualitatively very similar to the tree distribution obtained via Eq. (15) and also to the lineage distribution computed by solving Eq. (4).

As the fluctuations in size control increase (Fig. 1c), the snapshot distribution broadens and the birth size distributions become skewed. The distributions of forward and backward lineage are nearly indistinguishable, but the tree distribution leans towards significantly smaller sizes. For large division errors but small fluctuations in size control (Fig. 1d), the snapshot distribution also broadens, but the size distributions in forward (red), backward lineages (blue) and population trees (green) all differ significantly in this condition. Cells in backward lineages are typically bigger than in the population tree and these are bigger than in forward lineages.

We verify the accuracy of our predictions by agent-based simulations. To this end, we stochastically sample division sizes using Eqs. (18), calculate the interdivision times *τ*_*d*_ = ln(*s*_*d*_/*s*_0_)/*α*, determine the birth size after division from Eq. (3) and assign the remaining size to the other daughter. The distributions from the simulated population trees (dots in Fig. 1) are in excellent agreement with the theory (solid lines) in all cases.

The predicted differences in the birth sizes can be understood intuitively from the associated interdivision times of cells. In the presence of division errors, cells born larger have shorter division times, and they are hence over-represented in the population tree and the backward lineages when compared to forward lineages. On the contrary, when only cell size control fluctuates, cells dividing at smaller sizes are over-represented in the population tree. Curiously, our theory predicts that the latter phenomenon affects only the tree statistics but not the backward lineage. This is because the lineage statistics have the same division size controls *φ* (cf. Eqs. (4) and (16)). In the following, we provide a quantitative analysis of these effects based on the cell size moments.

## V. CONDITIONS FOR CELL SIZE HOMEOSTASIS

### A. Mean cell size and cell-to-cell variability

To obtain expressions for the cell-size moments in a forward lineage, we multiply Eq. (4) by *s*^*n*^, perform the integration and employ the binomial theorem. The result is

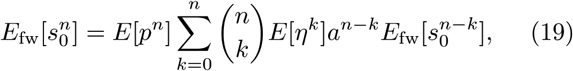

which depends on the moments *E*[*η*^*k*^] of the birth-size independent part of the division size (cf. Eq. (18)). Solving this relation for the *n*^*th*^ moment 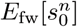 allows to express the cell size moments recursively in terms of lower order moments

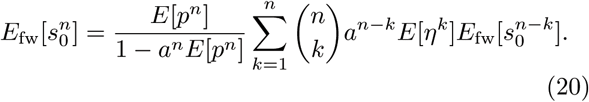

Using the above equation, we have explicit expressions for the mean cell size and its fluctuations. Similarly, the moments in backward lineages 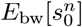 are obtained by replacing *E*[*p*^*n*^] with *E*_bw_[*p*] = 2*E*[*p*^*n*+1^] (cf. Eq. (17)), and the moments of the tree distribution follow from 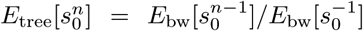 (cf. Eq. (15)). In summary, the mean size and its coefficient of variation 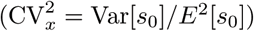 are given by

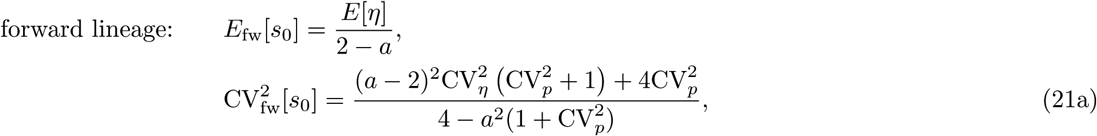

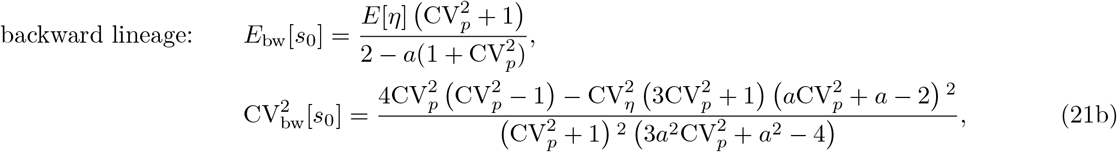

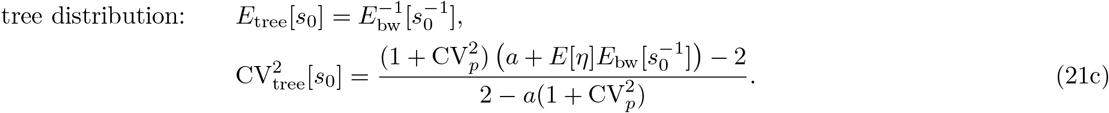

These expression are exact but provide exact analytical insights only to the forward and backward lineage statistics, because the tree mean cannot be obtained in closed form and has to be approximated. In what follows, we provide a discussion of quantitative differences between these different statistics, analyse their sensitivities to different noise sources, and compare our findings to experimental single-cell data.

#### 1. Quantitative differences of the cell-size statistics

We first compare the lineage statistics. Interestingly, the mean cell size in a forward lineage is independent of the division error CV_*p*_ while it increases with CV_*p*_ in backward lineages confirming our previous findings (Fig. 1d). We illustrate this dependence for the sizer (*a* = 0), adder (*a* = 1) and a timer-like size control (*a* = 1.5) in Fig. 2a. It is seen that the mean cell size in backward lineages is more sensitive to the cell size control parameter a. In Fig. 2b we compare the dependence of the coefficient of variation of size fluctuations for different size controls and division errors. Importantly, despite the sensitivity of the mean cell size to division errors, for sizer and adder the coefficient of variation is always smaller in the backward than in the forward lineage. It can be shown using Eqs. (21) that this holds whenever *a* ≤ 1.

**FIG. 2.**
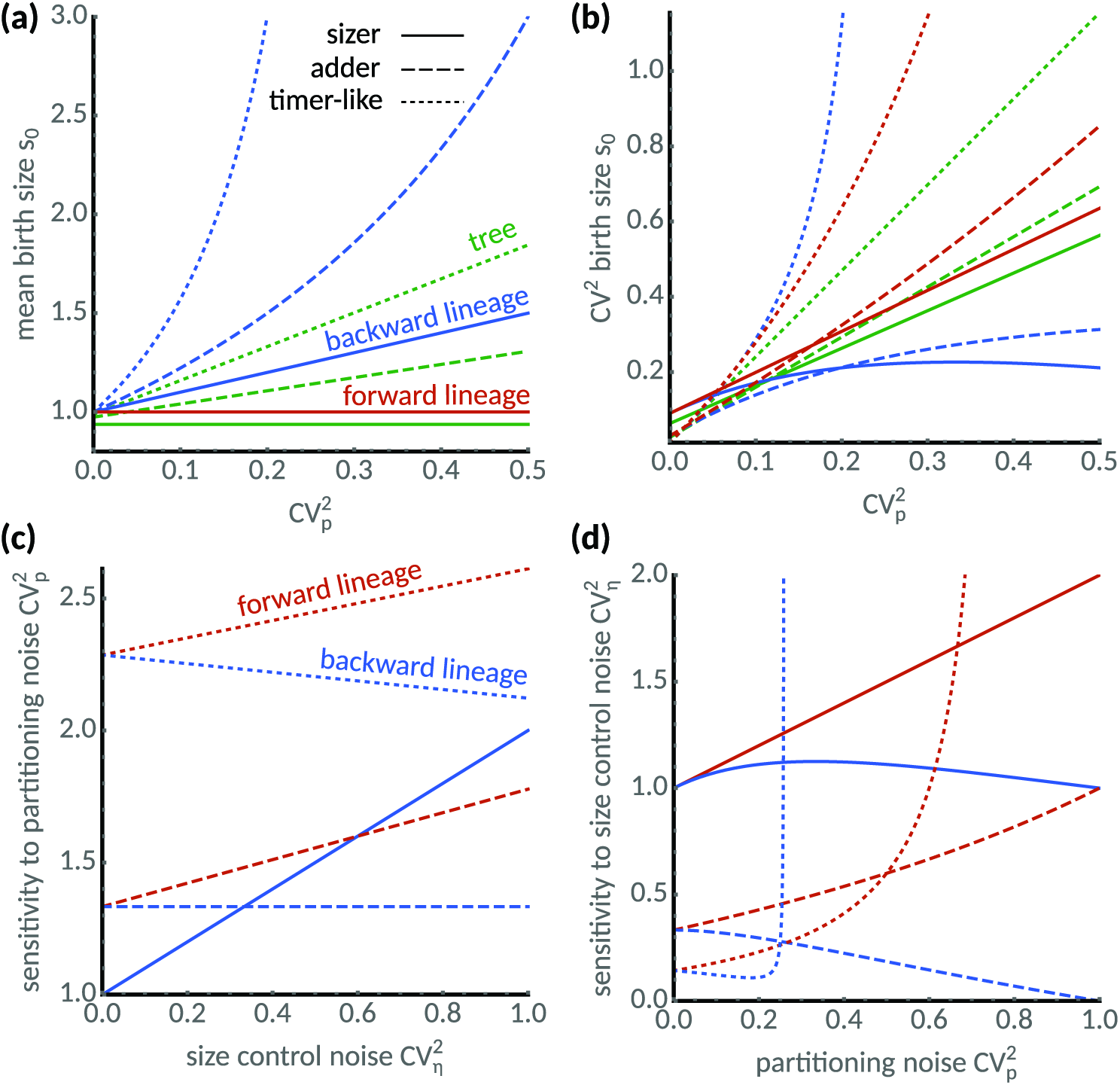
Cell size statistics and their sensitivities differ between single cells and populations. **(a)** Mean birth size increases with division errors in backward lineages (blue) but not in forward lineages (red). In the tree statistics, mean birth size can be smaller or larger than in forward lineages depending on partition errors. Predictions for various modes of cell size control are shown: sizer (*a* = 0, solid), adder (*a* = 1, dashed) and timer-like (*a* = 1.5, dotted). We chose *E*[*η*] = (2 − *a*), CV_*η*_ = 0.3 and skew[*η*] = 1 such that the mean cell size in forward lineages is independent of the cell size control, (b) Comparison of coefficients of variation of birth size in forward (blue), backward lineages (red) and lineage tree (green). For the sizer and adder mechanisms the size fluctuations in backward lineages and the tree statistic are always smaller than in forward lineages, **(c)** Sensitivity of the coefficient of variation of cell size to division errors in forward and backward lineages. Sensitivity increases with noise in cell size control in forward lineages (red lines) for all size controls. However, in backward lineages the sensitivity is increasing for the sizer, is constant for the adder and decreasing for the timer-like control, **(d)** Sensitivity of the coefficient of variation of cell size to noise in size control. For small division errors, sensitivity of the the sizer increases both in forward and backward lineages and the sensitivities of adder and timer-like controls display opposite behaviours in forward and backward lineages. Note that sensitivities diverge for the timer-like controls.

In the tree statistics, cells are expected to be smaller in the tree statistics than in backward lineages

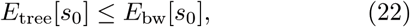

which follows using Jensen’s inequality in Eq. (21c). To compare the tree statistics with the forward statistics, we approximate the tree mean assuming small fluctuations by expanding Eq. (21c) about *E*_bw_[*s*_0_] and substituting the backward moments. A straight-forward calculation including also the skewness skew[*η*] of the size control distribution gives

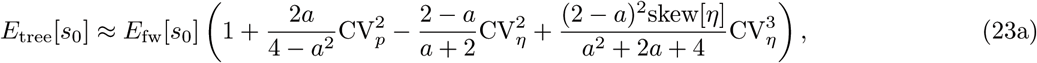

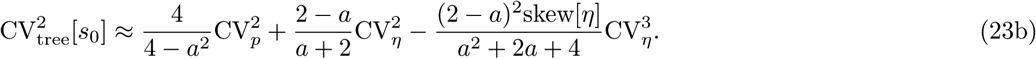

Note that by Eq. (8) division errors do not posses skew-ness and hence there is no corresponding term.

By inspecting the second and third term in the approximation of the tree mean, we observe that division errors increase the mean while errors in the size control decrease it relative to the forward lineage average. Interestingly, the tree mean of the sizer control (*a* = 0) decreases with size control errors but is insensitive to division errors, while a timer control (*a* = 2) amplifies division errors (Fig. 2a). For the adder control (*a* = 1), the tree mean can either be smaller or larger than the forward lineage average depending on the relative the relative size of the noise sources (Fig. 2a).

The differences in the coefficients of variation of the tree statistic and the forward lineage are determined by the skewness skew[*η*] of the size control distribution. This follows because the first two terms in Eq. (23b) equal the linearised coefficient of variation in forward lineages. Negative skewness hence increases the noise levels in the tree statistics, while positive skewness decreases it. The latter case is shown in Fig. 2b where we find that noise levels are smaller than forward lineage ones. Considering the fact that backward lineages also have smaller noise levels than forward lineages for *a* ≤ 1, this suggests a possible mechanism for noise reduction in populations.

#### 2. Sensitivity analysis

To investigate the different lineage statistics further, we compute the sensitivities of the coefficient of variation given in Eqs. (21) to the different noise sources

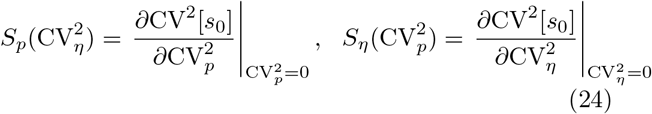

We find that the sensitivity to division errors 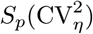 shown in Fig. 2c increases with noise in cell size control in forward lineages (red lines) for all size controls. However, in backward lineages, the sensitivity increases only for the sizer, it is constant for the adder and decreases for the timer control.

The sensitivity of size variation to noise in cell size control 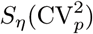) shown in Fig. 4d increases for the sizer mechanism with small division errors both in forward and backward lineages, but exhibits opposite dependencies for the adder and timer-like controls in the respective lineage statistics. We also observe that an instability occurs for timer-like mechanisms as described previously^22,24^. Interestingly, this instability occurs for smaller values of a in the backward lineages than those previously reported for forward lineages. This suggests that different conditions for cell size homeostasis apply to forward and backward lineages but also to the tree distribution. Before analysing these conditions in Sec. VB, we compare the predictions of our theory to single-cell data.

#### 3. Application to single-cell experiments

To validate the predictions of our theory, we analyse a recent dataset acquired using time-lapse microscopy of the mycobacterium *S. smegmatis*^8^. Its rod-shape allows us to use of cell length as a proxy for cell size. Fig. 3 summarises the lineage-weighted statistics of mean lengths and the corresponding coefficients of variation as introduced in Sec. II. We find that mean length is higher in the tree statistic and backward lineages than in forward lineages (Fig. 3a). Similarly, we observe smaller cell-to-variability in the tree and backward lineages (Fig. 3b). The noise reduction in the backward lineage noise is roughly 10%. These findings qualitatively confirm the predicted differences in the cell size statistics of populations.

**FIG. 3.**
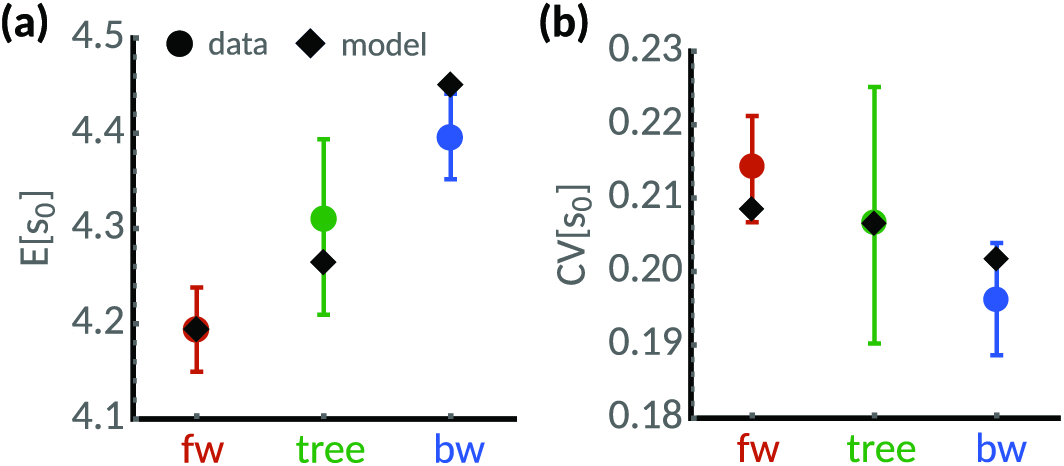
Cell-size statistics of the mycobacterium *S. smegmatis*. Analysis of the time-lapse microscopy data for cells grown in glycerol reported by^8^. (**a**) Mean cell length estimated using lineage-weighted statistics of backward lineages (blue) and tree statistics (green) is lower than in the forward lineage (red dots). (**b**) The corresponding coefficient of variation of cell size of backward lineages (blue) and tree statistics (green) is lower than in the forward lineage (red dots). Error bars denote 95%-confidence intervals obtained from bootstrapping. Diamonds denote model predictions for a adder sizer control (*a* = 1) using Eqs. (21a), (21b) and (23). Best-fit parameters obtained from nonlinear least-squares are *E*[*η*] = 4.19, CV_*p*_ = 0.17, CV_*η*_ = 0.10, skew[*η*] = 2.

To quantitatively compare these measurements with the proposed theory, we fit the expressions in Eqs. (21a), (21b) and (23) using nonlinear least squares. Because cells grown in glycerol follow the adder rule^8^, we fix *a* = 1. The best-fit (see caption of Fig. 3 for parameters) provides reasonable agreement with the measured quantities considering the experimental error bars. We expect that higher sample sizes and developed models of the size control accounting for the known asymmetric growth of old and new pole cells^8^ could improve the quantitative agreement.

### B. Moment conditions for cell size homeostasis

Next, we investigate the conditions for cell size ho-0 moeostasis on the basis of the existence of certain cell size moments. From the recursive relation in Eq. (20) we observe that if a certain moment is bounded, all lower order moments must be finite as well. Specifically, we can deduce the condition for the *n*^*th*^ moment in forward lineages to be bounded by inspecting the denominator in Eq. (20). Conditions for the backward lineages follow from substituting *E*[*p*^*n*^] → 2*E*[*p*^*n*+1^] and subsequently *n* → *n* − 1 for the tree distribution. In summary, the conditions for the existence of the *n*^*th*^ moment are:

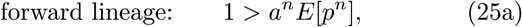

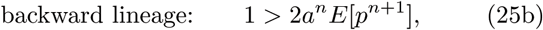

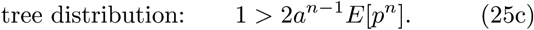

We make three important observations: (i) The above conditions are independent of the noise in size control but are identical only when the divisions are perfect *p* = 1/2; (ii) *a* < 1 guarantees all moments to be finite; and (iii) there exists a hierarchy between the different conditions. A finite moment of tree distribution implies the same moment to be finite in the forward lineage. A finite moment in the backward lineage implies the same moment to be finite in both the forward lineage and the tree distribution. This hierarchy follows because 2*a*^*n*^*E*[*p*^*n*+1^] ≤ 2*a*^*n*−1^*E*[*p*^*n*^] ≤ *a*^*n*^*E*[*p*^*n*^].

We will briefly discuss the marginal case of the adder rule (*a* = 1) because it models the size control of many microbes. Its moments are always finite in the forward lineage but not in the backward lineage and tree distribution. This extreme situation 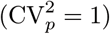 appears, however, only when one daughter inherits all of the mother’s size resulting in a non-growing cell and one that grows without bound. Such a situation has been observed in minicell-producing mutants of *E. coli*, but these cells also allow for divisions at mid-cell reinforcing their overall viability^30^. In the following, we investigate in more detail the conditions for finite mean and variance.

#### 1. Conditions for finite mean size and cell-to-cell variability

The conditions for the cell size distributions having finite mean values are found by letting *n* = 1 in Eqs. (25a), which leads to

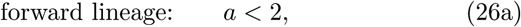

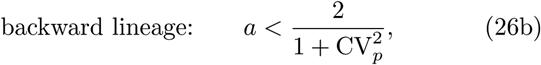

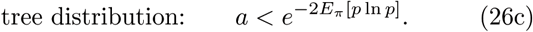

The first and second conditions are obtained from noting that *E*[*p*] = 1/2 and 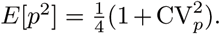. The third con dition is not immediately obvious since condition (25c) does not provide direct information about the mean. The result can be obtained by perturbing (25c) by *n*→ 1 + *∊* and requiring its leading order term to be positive.

Note that the condition for the forward lineage depends only on the cell size control. Interestingly this is not the case for the backward lineage and tree distributions, which depend intricately on the distribution of division errors. We summarise these findings in Fig. 4a, which shows the regions in parameter space *a* and 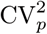 for which the first moments are finite. We observe that a finite mean in the backward lineage implies a finite mean in the other measures. Similarly, a finite mean of the tree distribution implies the same for the forward lineage but not for the backward lineage, as discussed in the previous section.

**FIG. 4.**
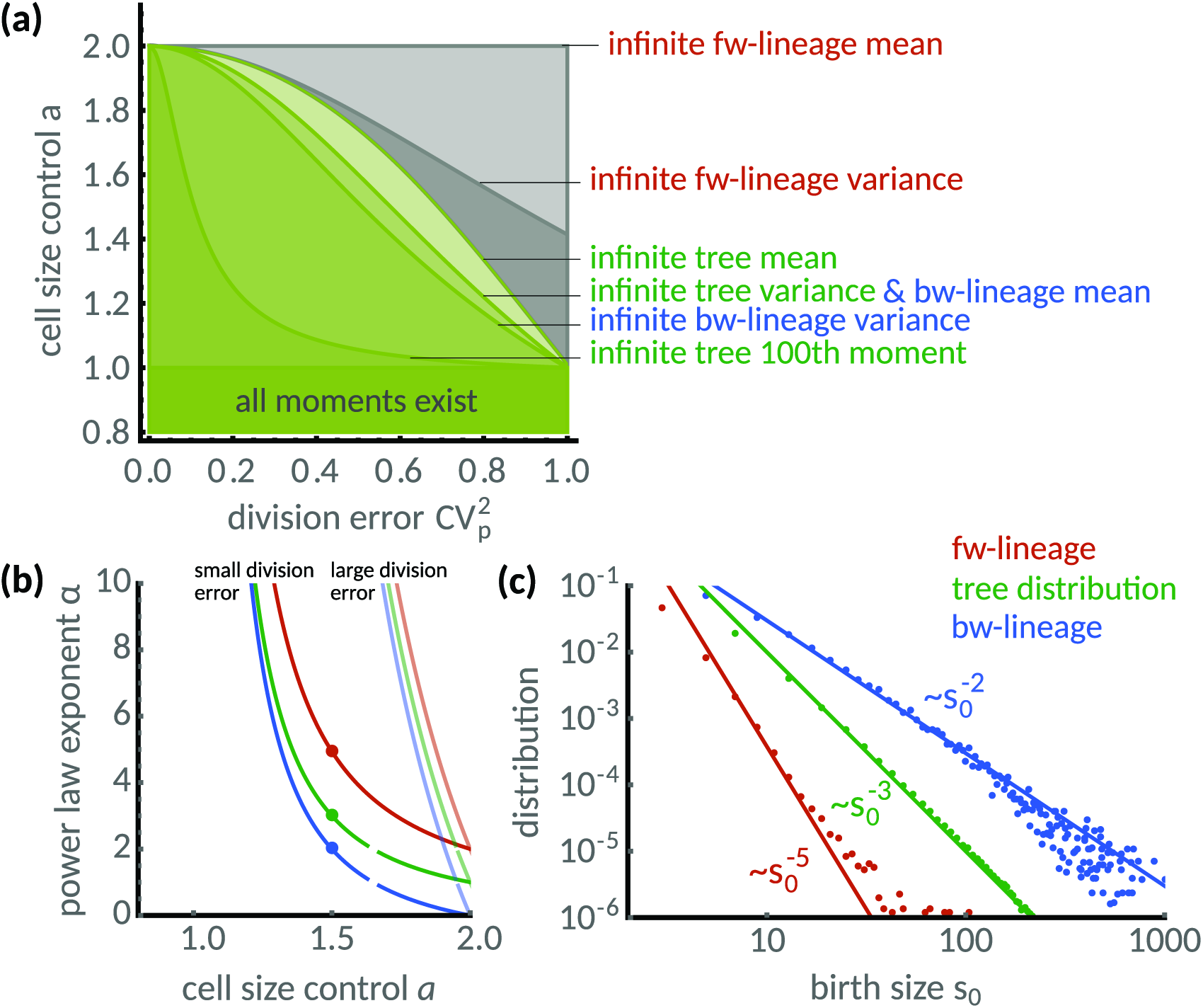
Distributions of timer-like mechanisms display different power laws in single cells and populations. **(a)** Regions in parameter space *a* (cell size control) and 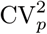 (division error) under which cell-size homeostasis is established in terms of first and second moments. Lines separate the regions above which forward lineages have infinite mean (light grey) and variance (dark grey), the population tree has infinite mean (very light green) and variance (light green) and similarly for the backward lineages (light and medium green). Also shown is the line above which the 100^*th*^ tree moment is infinite (dark green) computed for a symmetric beta distribution. Note that for *a* ≤ 1 all moments are finite. **(b)** Power law exponent of size distribution in dependence of the size control parameter *a*. The exponents are higher for low division errors (light colours, symmetric beta distribution with 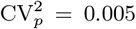) than for large division errors (full colours, 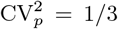) but are generally different for forward (red), backward lineages (blue) and tree distributions (green). **(c)** Comparison of power laws obtained from theory (*a* = 1.5, compare dots in (c)) show excellent agreement with stochastic simulations. The power law exponent is *α* ≈ 5 in forward lineages (first three moments exist). In contrast, *α* = 3 (finite mean but infinite variance) for the tree distribution and *α* = 2 (infinite mean and variance) for the backward lineage distribution.

Similarly, the conditions for existence of the second moments can be investigated, which hold identically for the coefficients of variation of cell size. The result is given by

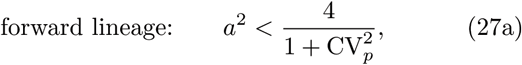

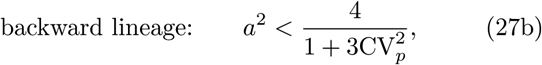

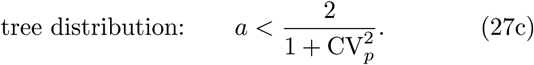

These conditions are obtained using 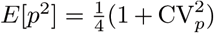 and noting that *π*(*p*) is symmetric about the mean and hence has no skewness leading to 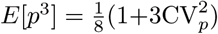 In Fig. 4a we illustrate the resulting regions where forward, backward lineages and tree variance are bounded.

### C. The distributions of timer-like controls display different power laws in single cells and populations

Divergent moments can be indicative of rare cell sizes and distributions with power law tails. To investigate the emergence of power laws in the different statistical measures, we assume that for large cell sizes the distributions follow 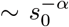 with power-law exponent *α*. The case *α* = ∞ corresponds to exponential tails while finite values of *α* indicate a power law. These can be deduced from the moments conditions because for a given power-law exponent *α* all moments of order *α*−1 are unbounded. Thus from the conditions (25), we find that the power-law exponents satisfy the following equations

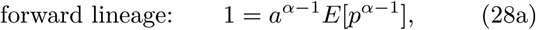

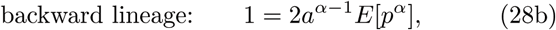

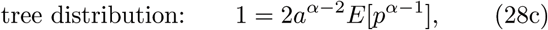

for *α* > 1. Because of the inequality given after Eq. (25), it follow that

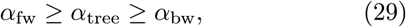

independently of the distribution of division errors. Note that the exponents are equal when divisions are perfect (*p* = 1/2).

However, it is difficult to give an explicit expression for the exponents without assuming a particular distribution *π*(*p*) of the division errors. In Fig. 4b, we study the dependence of the power-law exponent *α* on the cell size control parameter *a* for small and large division errors following a symmetric Beta-distribution. We find that the tails are exponential (*α* = ∞) whenever *a* < 1. For *a* > 1, however, the exponents of forward, backward lineages and tree distributions are similar only for small division errors and they deviate substantially for large division errors.

To study this case in more detail, we focus on the timer-like case *a* = 1.5 with large division errors. In Fig. 4c we compare the predicted power laws with stochastic simulations. The power law exponent of the forward lineage distribution is *α* ≈ 5 meaning that only the first 3 moments exist. In contrast, the tree distribution shows the exponent *α* = 3, which translates to finite mean cell size but unbounded variance. Interestingly, the backward lineage distribution has exponent *α* = 2, which does not admit a finite mean cell size. In all cases, the simulations (dots) are in excellent agreement with the theoretical exponents (lines).

## VI. DISCUSSION

Cell size has been traditionally quantified by snapshots, but recent advances in time-lapse microscopy allow to track the growth of hundreds to thousands of cells in a microcolony. These techniques allow to construct entire lineage trees and resolve genealogies and thus offer various statistics with which to quantify individual cell sizes. We presented a unified modelling framework to quantify and compare size distributions in these populations.

We demonstrated that cells in populations exhibit increased cell size, often exhibit lower cell-to-cell variability and display different sensitivities to division errors and cell cycle noise compared to cells in isolation. Specifically, we found that positive skewness in the size control distribution decreases cell-to-cell variability in the population tree. In fact, many bacteria implementing the adder control such as *E. coli*, *B. subtilis*^6^ and *S. smegmatis*^8^, show positively skewed length increments. We also observed that whenever *a* ≤ 1, the coefficient of variation of cell size in backward lineages is smaller than in forward lineages. Since backward lineages represent typical lineages in a population tree, we speculate that populations implementing adder or sizer controls, which are ubiquitously found in microbes, could exploit these differences to narrow their size distributions.

A general requirement for size homeostasis, regardless of the model, is that cells born small grow proportionally more than cells born with larger sizes^9^. These larger cells proliferate quicker and are therefore over-represented in populations, which explains the discrepancies we found between different size distributions, their associated moments and power laws in the presence of division errors. We hence expect our conclusions to hold also for developed models of cell size control. We found that these differences are particularly sensitive to division errors, which can be extremely heterogeneous when cells respond to stress^?^. Conversely, we found that cells dividing at smaller sizes are over-represented in the population tree when division size is highly variable. We anticipate that this effect will be important when cell size control is modulated by time-dependent environments^31^.

Finally, we elucidated that isolated cells tracked over many generations and cells in populations exhibit different sensitivities to division errors and noise in cell cycle control. Competing experimental devices such as mother machines and agar pads may thus give qualitatively different conclusions about a cell’s ability to control its size and the resulting population heterogeneity. Similar effects have been described for stochastic gene expression^32^. However, our findings raise a more fundamental question: what type of size distribution cells may have evolved to control?

## ACKNOWLEDGEMENTS

PT thanks Miles Priestman for sharing the raw data to produce Fig. 3.

